# Neonatal Zika virus infection causes epileptic seizures, viral persistence and long-term behavioral impairment

**DOI:** 10.1101/195172

**Authors:** Isis N. O. Souza, Paula S. Frost, Julia V. França, Jéssica Nascimento-Viana, Rômulo L. S. Neris, Leandro Freitas, Daniel J. L. L. Pinheiro, Gilda Neves, Leila Chimelli, Fernanda G. De Felice, Ésper A. Cavalheiro, Sergio T. Ferreira, Andrea T. Da Poian, Iranaia Assunção-Miranda, Claudia P. Figueiredo, Julia R. Clarke

**Affiliations:** School of Pharmacy, Federal University of Rio de Janeiro, Rio de Janeiro, RJ 21944-590, Brazil; Institute of Biomedical Sciences, Federal University of Rio de Janeiro, Rio de Janeiro, RJ 21944-590, Brazil; Institute of Microbiology Paulo de Goes, Federal University of Rio de Janeiro, Rio de Janeiro, RJ 21944-590, Brazil; Department of Neurology and Neurosurgery, Escola Paulista de Medicina, Federal University of São Paulo, São Paulo, SP 04023-062 Brazil; Laboratory of Neuropathology, State Institute of Brain Paulo Niemeyer, Rio de Janeiro, Brazil; Institute of Medical Biochemistry Leopoldo de Meis, Federal University of Rio de Janeiro, Rio de Janeiro, RJ 21944-590, Brazil; Centre for Neuroscience Studies, Department of Biomedical and Molecular Sciences, Queen’s University, Kingston, ON, Canada; Institute of Biophysics Carlos Chagas Filho, Federal University of Rio de Janeiro, Rio de Janeiro, RJ 21944-590, Brazil

**Keywords:** Zika virus, perinatal infection, seizures, epilepsy, neuroinflammation, necrosis

## Abstract

A causal relationship between congenital Zika virus (ZIKV) exposure and microcephaly and other neurological disorders have been established, but long-term consequences of infection are still unknown. We evaluated acute and late neuropathological and behavioral consequences of ZIKV infection in a neonatal immunocompetent mouse model. ZIKV showed brain tropism, causing post-natal microcephaly and several behavioral dysfunctions. During the acute phase of infection, mice developed very frequent epileptic seizures, which are consistently reduced by TNF-α neutralization. Although adult animals recover from seizures, they become more susceptible to chemically-induced crises. Intriguingly, the virus remained actively replicating in adult animals, which show persistent necrosis and calcifications in the mice brain. Altogether the results reveal late consequences of neonatal ZIKV exposure and suggest the early inhibition of neuroinflammation as a potential treatment.

## Introduction

Zika virus (ZIKV) is an arbovirus responsible for a major outbreak in the Americas in 2015 *(1)*. Similar to other flaviviruses, *Aedes* mosquitoes are the main vector for ZIKV, but transmission through sexual contact and blood transfusion have also been reported *(2–4)*. Clinical and experimental studies have shown that ZIKV is neurotropic and can be especially harmful to the immature nervous system *(5)*. ZIKV has been isolated from brains of newborns with congenital microcephaly, and a causal relationship between *in utero* viral exposure and severe neurological malformations has been firmly established *(6)*.

While extensive research has focused on understanding the mechanisms of ZIKV-induced congenital microcephaly, the most dramatic effect reported in ZIKV-infected newborns, it is now known that only 1-13% of exposed fetuses show physical malformations at birth *(7)*. Moreover, a spectrum of clinical manifestations appears months to years after birth, both in normo- and microcephalic babies exposed to ZIKV during gestation *(8, 9)*. For example, babies infected during pregnancy and born with normal head sizes can develop postnatal microcephaly, suggesting that ZIKV has consequences beyond the gestational period *(10, 11)*. Further, hemiparesis, dyskinesia/dystonia, dysphagia, persistence of primate reflexes, developmental delays and motor impairment have been reported in normocephalic babies with laboratory evidence of congenital ZIKV exposure *(12–14)*.

Seizures are recurrent events in infants who developed ZIKV-induced microcephaly, and reports suggest that 60% of normocephalic infants exposed to ZIKV *in utero* suffer similar symptoms *(11, 15, 16)*. Cortical dysplasia induced by ZIKV has also been described in babies born with normal head sizes, a finding often underdiagnosed and that can itself be associated to seizures *(11, 17)*. Thus, it is now clear that focusing only on microcephaly and other physical malformations will lead to an underestimation of the true magnitude of the health consequences of the ZIKV epidemics.

Experimental models for investigating the effects of ZIKV on the developing brain have mainly focused on transgenic immunodeficient mice *(18–21)* and *in vitro* organoid cultures *(5)*. Few studies have described the effects of ZIKV infection in the developing brain of immunocompetent rodents *(22–24)*, but the late-life consequences of early life infection have not been addressed so far. Here, we infected newborn immunocompetent mice systemically with a Brazilian ZIKV strain and performed a long-term follow-up of the behavioral, neuropathological and molecular consequences of infection. The results showed that ZIKV reached and replicated in the developing brain, inducing weight loss, mortality and up-regulation of several neuroinflammation markers. Neonatal ZIKV infection further caused brain atrophy, severe motor dysfunction and episodes of seizures in young mice. Most strikingly, we demonstrated that although the animals exposed to ZIKV in the neonatal period recover from the seizures in adult life, they show an increased susceptibility to chemically-induced seizures, together with other severe behavioral dysfunctions such as cognitive and social impairments. Our findings warn of the need for special attention to a potential burden of neuropathological complications in children and adolescents after congenital exposure to ZIKV.

## Methods

***Virus*.** ZIKV was isolated from a febrile patient in the state of Pernambuco, Brazil (gene bank ref. number KX197192). The stocks used in the experiments were produced and tittered as previously described in Coelho et al., 2017 *(25)*. As a control, the same volume of virus-free conditioned medium of C6/36 cells was used (mock). In some experiments, a second control group comprising inactivated ZIKV was used. Inactivated ZIKV was obtained by direct exposure of the virus solution for 30 min to UV-light. Successful inactivation was confirmed by subsequent plaque assay.

***Animals and neonatal infection*.** All procedures used in the present study followed the “Principles of Laboratory Animal Care” (US National Institutes of Health) and were approved by the Institutional Animal Care and Use Committee of the Federal University of Rio de Janeiro (protocol #052/2017). Naïve 10 week-old female Swiss mice were obtained from our facilities and mated for two weeks (four females per male). After this period, dams were housed individually until delivery. On the third day after birth (P3), each litter was left with five to ten pups, with equal numbers of males and females whenever possible, and were infected subcutaneously (s.c.) with 30 μL of ZIKV (10^6^ PFU) or the same volume of mock medium. All littermates received the same treatment in order to avoid cross-contamination. Mice showing any signs of reflux or hemorrhage (∼ 5% of animals throughout our study) were excluded from further analyses. In some experiments, a control group of mice was injected with UV-inactivated ZIKV, as described above. Pups were evaluated daily for mortality, and body weight was measured every three days from the day of infection to 66 days post-infection (dpi).

In experiments aimed at evaluating adult behavior, mice were weaned at post-natal day 21 (P21) and housed with same-sex littermates (2-5 mice per cage). Mice were housed in polypropylene cages and maintained at 25^o^C with controlled humidity, under a 12 h light/dark cycle and *ad libitum* access to water and chow. In order to rule out any effect of early maternal separation on behavior, independent groups of mice were used for pre-weaning experiments - evaluation of post-natal reflexes and seizure recordings.

For determination of viral replication and cytokine mRNA levels, mock-, ZIKV- or inactivated ZIKV-injected mice were killed by decapitation at different times post infection and brains were rapidly removed and frozen in liquid nitrogen until RNA isolation for qPCR analyses.

***Neonatal reflexes*.** Temporal development of grasping reflex, negative geotaxis and righting reflex was evaluated in newborn mice from the day of infection to 13 dpi. Impaired/delayed reflexes or their complete absence in newborns usually indicates abnormalities in neurodevelopment. The righting reflex is the ability of mice to rapidly return to ventral decubitus when placed in dorsal decubitus on a bench. Two randomly chosen pups from each litter were placed on their backs over a flat surface and were given one minute to complete the task. The grasping reflex is assessed by placing a blunt object on the paw of newborn pups. For this procedure, mice were held by the scruff of their necks and each front paw was individually stroked with the blunt end of a small paperclip. Scores were given as such: 2 if the reflex is present in both front paws; 1 if the reflex is present in only one front paw; 0 if the reflex is absent in both front paws. The negative geotaxis test evaluates not only motor coordination but also the quality of vestibular cue for perception of gravity. Mice were placed on a metallic grid inclined 30° with their heads facing downwards and were given one minute to turn 180°, finishing the task with body and head facing upwards. Mice that fell from the apparatus were retested.

### Behavioral tests

***Hindlimb suspension*.** At 9 dpi, animals were individually removed from their homecages and placed inside a 50mL laboratory tube padded with cotton balls, facing the interior of the tube, suspended by the hind limbs. Time to fall inside the tube was measured as an assessment of muscle strength and general neuromuscular function.

***Pole test*.** Mice at 15 to 21 dpi were placed head-up at the top of a vertical wooden pole (height: 50 cm; diameter: 1.5 cm) covered by rough clothing. The base of the pole was placed in a mouse homecage containing clean sawdust. When placed at the top of the pole, animals orient themselves downwards and descend the length of the pole back into the cage. Animals underwent 2 days of training with five trials each to learn the task. On the third day, a test session was performed, and animals were again placed at the top of the pole and the time to orient downwards (time to turn) as well as the total time to descend the pole (time to descend) were measured; the three best performances of each animal were selected and used for the statistical analysis, as previously described *(26, 27)*.

***Rotarod*.** This test was performed in the rotarod apparatus (Insight LTDA, Brazil) at ∼90 dpi. For habituation, mice were individually placed in the apparatus floor for 3 minutes and then on the aluminum cylinder without rotation for another 2 minutes. Afterwards, animals were submitted to three test sessions (∼1 hour intervals) when each mouse was placed on the aluminum cylinder at a speed of 5 RPM, and was accelerated until 35 RPM during 5 minutes. Latency to fall the cylinder was measured, and the mean of the sessions was used. Mice that fell before 15 seconds were retested.

***Open field test***. Mice at ∼90 dpi were individually placed at the center of an arena (30 x 30 x 45 cm). Total distance traveled in the apparatus was recorded for 5 minutes by ANY-maze software (Stoelting Company). The arena was thoroughly cleaned with 70% ethanol in between trials to eliminate olfactory cues and illuminated with an indirect source of light (∼100 lux).

***Novel object recognition test*.** The test was carried out in the same day and in the same box used for the open field test. Test objects were made of plastic and had different shapes, colors, sizes and textures. During behavioral sessions, objects were fixed to the box using tape to prevent displacement caused by exploratory activity of the animals. Preliminary tests showed that none of the objects used in the experiments evoked innate preference. Before training, each animal was submitted to a 5 min-long habituation session, in which it was allowed to freely explore the empty arena. Training consisted in a 5 min-long session during which animals were placed at the center of the arena in the presence of two identical objects. The time spent exploring each object was recorded by a trained researcher. Sniffing and touching the object were considered exploratory behaviors. The arena and objects were cleaned thoroughly with 70% ethanol in between trials to eliminate olfactory cues. Ninety minutes after training, animals were again placed in the arena for the test session, in which one of the objects used in the training session was replaced by a new one. Again, the time spent exploring familiar and novel objects was measured. The results were expressed as percentage of time exploring each object during the training or test sessions. Animals that recognize the familiar object as such (i.e., learn the task) explore the novel object > 50% of the total time *(28)*.

***Social approach test*.** The social approach test was performed at ∼100 dpi in a three-chamber apparatus, which consisted in a rectangular transparent acrylic box (60 cm x 45 cm x 30 cm) with two walls dividing the box in three equal chambers of 20 cm x 45 cm x 30 cm each. Cylindrical aluminum cages of 8 cm of diameter (9.5 cm height) were used to contain a stranger mouse (a mouse with which the test mice have never had any contact before). Sociability was assessed by placing one empty cage in one lateral chamber and one cage containing the stranger mouse in the opposite chamber. The test mouse was kept in the middle chamber for 5 minutes, after which the walls were removed and the mouse was allowed to fully explore the apparatus for 10 minutes. Time exploring each cage was manually recorded by a trained researcher.

***Seizure evaluation*.** Dams and their respective pups were removed from their home cages at different times of the light cycle selected at random and placed in a box measuring 41 cm x 34 cm x 18 cm containing clean sawdust, for one hour, daily from 9 to 15, 18 and 100 dpi. Male and female pups were individually identified with permanent ink in their backs, and high-resolution recording was performed using a Nikon D3300 camera fixed on the ceiling of the room. Videos were analyzed by an experienced researcher who identified animals showing any degree of motor seizures *(29)*. The percentage of animals showing seizures per litter during the one-hour session was determined.

***Pharmacological treatments*.** Swiss mice were infected with 10^6^ PFU of ZIKV on postnatal day 3 (P3) and randomly assigned into one of the following groups: PBS; infliximab (20 µg/day) or N-acetylcysteine (NAC; 50 µg/kg/day) *(30, 31)*. Treatment was given once a day intraperitoneally (ip) from the day of infection to 12 dpi. Development of seizures was assessed at 12 dpi in an one hour-long recording sessions as described above.

***Electrographic recordings*.** Mice at 10 to 12 dpi and ∼100 dpi were anesthetized (Isoflurane) and implanted with gold plated electrodes (1.0 mm length; 2 in the right and 2 in the left cortical surface; Supplementary Figure 1A). This allowed right and left cortical activity could be independently analyzed. Three days after surgery, mice were individually placed in a Plexiglas box measuring 41 cm x 34 cm x 18 cm. The external tip of the electrodes was then plugged to a connector linked to isolated wires that could reach the amplifier (FE136 Animal Bio Amp – AD Instruments). Signals were then digitalized (PowerLab 8/35 - ADInstruments) and saved in LabChart 8 (AD Instruments). Each animal was continuously recorded for 2 h. After recordings, animals were returned to their homecages. The same animals were recorded for up to 3 consecutive days.

**Figure 1.**
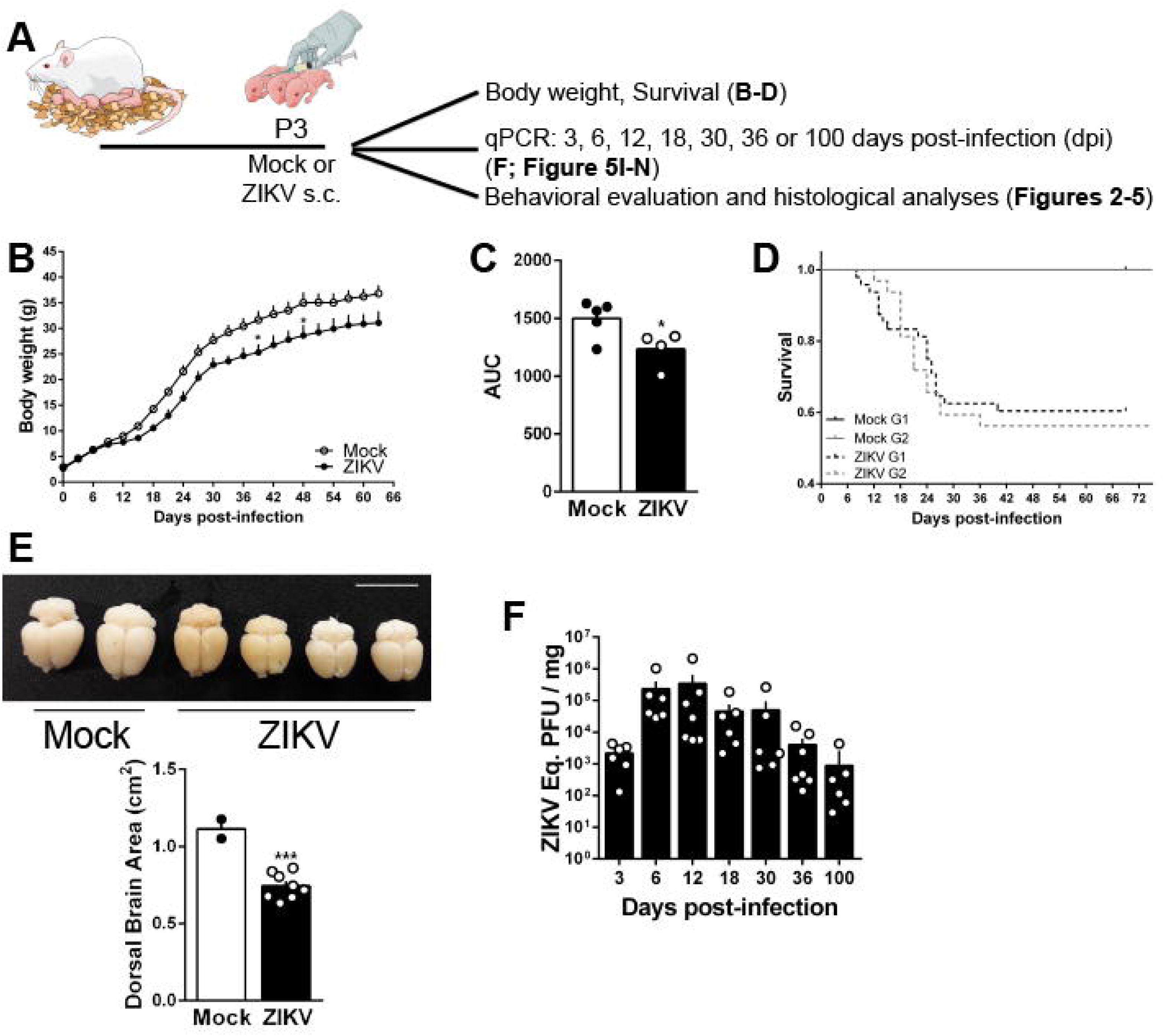
ZIKV replicates in neonatal mouse brain and induces weight loss, mortality and brain atrophy. (A) Schematic representation showing the experimental design, in which Swiss mice received a subcutaneous (s.c.) injection of 10^6^ PFU ZIKV or mock medium at post-natal day 3 (P3). (B) Body weight was measured every three days following ZIKV infection up to 66 days post-infection (dpi) and (C) area under the curve from body weight curves was calculated. (D) Representative survival curve from two independent experiments (G1 = group 1; G2 = group 2) (n = 4-5 litters/group in each experiment). (E) Representative images of brains from mock and ZIKV-injected mice perfused at 23 dpi. Bar graphs represent dorsal brain area quantification. (F) At the indicated days post infection, brains of mice were dissected and processed for determination of ZIKV mRNA by qPCR. In **B**: *p<0.05, two-way ANOVA followed by Sidak. In **C** and **E**: *p = 0.0413, ***p = 0.0006, Student’s *t* test.

***Pentylenetetrazol (PTZ)-induced seizures*.** Pentylenetetrazol (Sigma-Aldrich) was dissolved in saline to a final concentration of 1% and frozen until the day of experiment. Adult mice (∼110 dpi) received pentylenetetrazol (50 mg/kg, ip) and were immediately placed in empty 3 L glass beakers. Latency to first seizure and number of seizures were evaluated by an experienced researcher during the following 20 minutes.

***Tissue collection and preparation*.** Mice were killed by decapitation at 3, 6, 12, 18, 30, 36 or 100 dpi for qPCR and at 23 and 100 dpi for histological and immunohistochemical analyses. For mRNA extraction, brains were rapidly removed, frozen in liquid nitrogen and maintained at -80^o^C until the moment of extraction. For histology and immunohistochemistry, mice were deeply anesthetized with ketamine (80 mg/kg) and xylazine (10 mg/kg) before transcardiac perfusion with cold saline solution followed by fresh ice-cold 4% formaldehyde. Brains were removed, post-fixed for 24 h in 4% formaldehyde, and embedded in paraffin after dehydration and diaphanization. Macroscopic atrophy of the brains was assessed by the determination of total dorsal brain area, using the Image J software.

***Quantification of reactive oxygen species*.** Brains of mock and ZIKV-infected mice were dissected at 12 dpi and gently homogenized in cold DMEM (Invitrogen; 1:10 w/v) using a glass tissue grinder. Reactive oxygen species (ROS) was determined in each sample by adding 2 μM of 5-(and-6)-chloromethyl-2’,7’-dichlorodihydrofluorescein diacetate acetyl ester (CM-H2DCFDA - Invitrogen) to 200 μl of a 1:1 mixture of homogenate/DMEM, followed by incubation for 30 minutes in the dark at 37°C. DCF fluorescence was quantified using a fluorescence microplate reader (VICTOR(^TM^) X3 – PerkinElmer), at 492-495/517-527 nm, as recommended by manufacturers.

***RNA extraction and qPCR*.** Brains were homogenized to a concentration of 0.5 mg tissue/mL in DMEM, and 200 μL of the homogenate were used for RNA extraction with Trizol (Invitrogen) according the manufacturer’s instructions. Purity and integrity of RNA were determined by the 260/280 and 260/230 nm absorbance ratios. Only preparations with ratios >1.7 and no signs of RNA degradation were used. One μg of total RNA was treated with DNAse I (ThermoFisher Scientific Inc) according to manufacturer’s recommendations before cDNA synthesis using the High-Capacity cDNA Reverse Transcription Kit (ThermoFisher Scientific Inc). For negative strand quantification cDNA was synthesized using 2 pM of the primer 835 as previously described *(32)*. Analyses were carried out on an Applied Biosystems 7500 RT–PCR system using the TaqMan Mix (ThermoFisher Scientific Inc). Primers for ZIKV were used as described by Lanciotti (2008): forward, 5’-CCGCTGCCCAACACAAG-30; reverse, 5’-CCACTAACGTTCTTTTGCAGACAT-3’; probe, 5’-/56- FAM/AGCCTACCT/ZEN/TGACAAGCAATCAGACACTCAA/3IABkFQ/-3’ (Integrated DNA Technologies). Cycle threshold (Ct) values were used to calculate the equivalence of log_10_ PFU/mg tissue after conversion using a standard-curve with serial 10-fold dilutions of a ZIKV stock sample. For cytokine and iNOS expression, qPCR analyses were performed using the Power SYBR kit (Applied Biosystems; Foster City, CA). Actin was used as an endogenous control. Primer sequences were the following: IL-6 Forward: 5’-TTC TTG GGA CTG ATG CTG GTG-3’ and Reverse: 5’-CAG AAT TGC CAT TGC ACA ACT C-3’; TNF-α Forward: 5’-CCC TCA CAC TCA GAT CAT CTT CT-3’ and Reverse: 5’-GCT ACG ACG TGG GCT ACA G-3’; IL-1β: GTA ATG AAA GAC GGC ACA CC and Reverse: ATT AGA AAC AGT CCA GCC CA; iNOS Forward: GTT CTC AGC CCA ACA ATA CAA GA and Reverse: GTG GAC GGG TCG ATG TCA C; KC: Fwd: CAC CTC AAG AAC ATC CAG AGC and Reverse: AGG TGC CAT CAG AGC AGT CT; Actin Forward: 5’-TGT GAC GTT GAC ATC CGT AAA-3’ and Reverse: 5’-GTA CTT GCG CTC AGG AGG AG-3’.

***Histological and immunohistochemistry analyses*.** Paraffin-embedded brain tissue sections (3-5 μm) were immersed in xylene for 10 minutes, rehydrated by incubation in absolute ethanol followed by 95% and 70% solutions of ethanol in water. For general histology, sections were stained by haematoxylin-eosin and imaged by light microscopy. For immunohistochemistry, slides were incubated with 3% H_2_O_2_ in methanol for inactivation of endogenous peroxidase. Antigens were reactivated by treatment with 0.01 M citrate buffer for 40 min at 95°C. Slides were washed in PBS and incubated with primary antibodies (Rabbit anti-Iba-1, Abcam, 1:200; rabbit anti-GFAP, DAKO, 1:500) for 12-16 hours at 2-8°C. After washing with PBS, slides were incubated with biotinylated secondary antibodies (Vector, 1:500) for 1 hour at room temperature, washed twice with PBS and incubated with streptavidin-biotin-peroxidase (Vector) for 30 min. The peroxidase reaction was visualized with 3,3’-diaminobenzidine (DAB) substrate for 1 to 5 min or until a brown precipitate could be observed. Identical conditions and reaction times were used for slides from different animals (run in parallel) to allow comparison between immunoreactivity densities. Reaction was stopped by immersion of slides in distilled water. Counter-staining was performed with Harris hematoxylin. Slides were washed in running water, dehydrated in alcohol, cleared in xylene, mounted in resinous medium and examined with light microscopy using a Sight DS-5M-L1 digital camera (Nikon, Melville, NY) connected to an Eclipse 50i light microscope (Nikon) at different magnifications.

***Statistical analysis*.** Statistical analysis was performed using GraphPad Prism 6.01 (GraphPad, USA). Data are reported as means ± S.E.M. Body weight curves were compared using two-way ANOVA followed by Sidak. In the object recognition test and social approach test were used one-sample *t* test compared to the fixed value of 50; All other comparisons used unpaired Student’s t test or one-way ANOVA followed by Tukey when adequate. Significance threshold was set at p ≤ 0.05.

## Results

### ZIKV replicates in neonatal mouse brain and induces weight loss, mortality and brain atrophy

In order to determine the progression of infection by ZIKV in the developing brain and its long-term consequences, immunocompetent Swiss mice were infected subcutaneously on post-natal day 3 with 10^6^ PFU of a Brazilian ZIKV isolate or an equivalent volume of mock medium (Fig. 1A). This period of rodent brain development and synaptic sprouting correlates with the last trimester of gestation in humans, hence our model simulates fetuses exposed during late gestation *(33)*. Since persistent weight loss has been associated with the severity of ZIKV infection in mice *(22, 24)*, mock- or virus-exposed pups were weighed every three days until 66 dpi. From 15 dpi onwards, body weight of ZIKV-infected mice was significantly lower than that observed for mock-injected animals (Fig. 1B, C). Pups were daily assessed concerning general appearance and mortality. Whereas all mock-infected pups survived throughout the duration of the experiment (up to 69 dpi), a 40% mortality was observed in the ZIKV-infected groups, beginning at 9 dpi and ceasing at 37 dpi (Fig. 1D). Two representative survival curves obtained in independent experiments are shown in Figure 1D, revealing a critical period of mortality in ZIKV-infected mice (between 12-30 dpi). We next investigated whether ZIKV replication was associated with macroscopic alterations in the brains of mice which survived the acute phase of infection. Since post-natal microcephaly has already been reported in fetuses exposed to ZIKV and born with normal head perimeter *(10, 11)* we determined whether infection in mice at post-natal day 3 induced alterations in brain size compared to mock-treated animals. We found a marked brain atrophy, as demonstrated by the reduced dorsal brain area in ZIKV-infected mice 23 dpi compared to mock-injected mice (Fig. 1E).

To determine whether peripherally-administered ZIKV reached and replicated in the developing brain, a distinct group of animals was infected as above and their brains were collected at 3, 6, 12, 18, 30, 36 or 100 dpi. ZIKV RNA was detected in the brain as early as 3 dpi and reached a peak between 6-12 dpi (Fig. 1F). This appears to be an important time window in experimental infection with ZIKV, as it comprises the peak of brain viral load, the onset of weight loss and the beginning of mortality. After this period, the level of viral RNA in the brain decreased progressively but remained detectable in 100 dpi adult mice (Fig. 1F). To determine whether viral RNA found at this late timepoint corresponded to actively replicating virus or residual fragments of viral RNA, we performed qPCR for the negative strand of ZIKV RNA. Interestingly, significant levels of the negative RNA strand were still detectable 100 days after infection (Suppl. Fig. 2A), suggesting that active replication is still occurring in brains of adult mice submitted to infection at post-natal day 3.

### Neonatal ZIKV infection induces seizures in young mice and increases susceptibility to chemically-induced seizures in adult mice

Human perinatal infections have been associated to seizure events throughout life *(34–36)*. In the specific case of ZIKV exposure, clinical/epidemiological studies have shown that both microcephalic and normocephalic babies exhibit spontaneous seizures at different intervals after birth *(10, 11, 16, 37, 38)*. Consistent with clinical findings, ZIKV-infected mice showed spontaneous seizures beginning around 9 dpi. To determine the percentage of ZIKV-infected mice that showed episodes of seizures, we video recorded the animals in their cages for 1 hour daily at random times of the light cycle. On average, 65% of the mice in each litter exhibited seizures at 9 dpi, and this increased to 88% of the pups at 12 dpi (Fig. 2A), coinciding with the peak of brain viral replication in the brain (see Fig. 1F). The percentage of mice presenting seizures remained high up to 15 dpi, decreasing significantly (to about 30%) by 18 dpi, to become undetectable in 100 dpi adult animals (Fig. 2A). No seizures were detected in mock-injected mice, and only two mice injected with UV-inactivated ZIKV (iZIKV) showed seizures in the recording session performed at 9 dpi, but not on subsequent sessions (Suppl. Fig. 1B).

**Figure 2.**
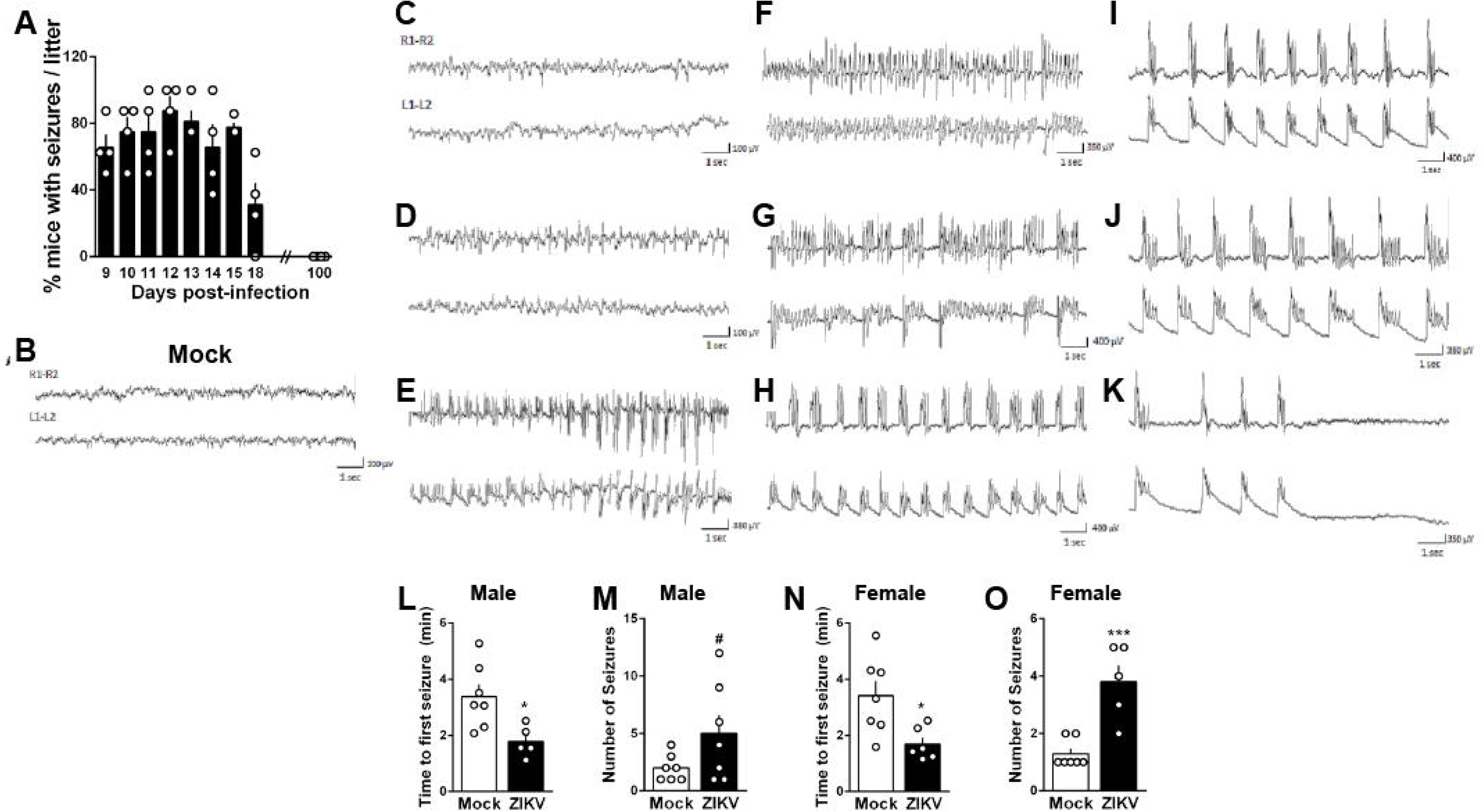
Neonatal ZIKV infection induces seizures in young mice and increases susceptibility to chemically-induced seizures in adult mice. Mice were infected by subcutaneous injection of 10^6^ PFU ZIKV or mock medium at post-natal day 3. (A) Percentage of animals with seizures in each litter was obtained from daily one hour-long recording sessions performed from 9 to 15 dpi, and at 18 and 100 dpi in ZIKV-infected mice. (B) Representative electroencephalographic recordings obtained from one mock-injected mice at 12 dpi. (C-K) Representative electroencephalographic recordings performed in brains of ZIKV-infected mice at 12 dpi show an initial period of apparently normal brain activity (C), followed by spiking activity more evident in the right hemisphere (D), which after 1-2 minutes evolved to a typical epileptiform activity with different graphoelements (fast spikes/fast sharp/polispike or polisharp-waves) (E-G). Interspersed occurrence of depressed activity with the epileptic activity was progressively observed (H-J) and in the following minutes culminated with the interruption of the epileptiform activity (K). n = 4 ZIKV-infected and 2 mock mice. (L-O) Time to first seizure and number of seizures for male (L, M) and female (N, O) mice after an ip injection of pentylenetetrazol at ∼110 dpi. Data are expressed as mean ± S.E.M. In **L:** *p = 0.0155, in **M**: **#**p = 0.0962, in **N**: *p = 0.0154, in **O**: ***p = 0. 0008. Student’s *t* test.

To perform an electrophysiological assessment of epileptiform activity induced by ZIKV, mice were submitted to surgery for implantation of cortical electrodes at the peak of viral replication and of seizures (Suppl. Fig. 1A). ZIKV-infected mice recorded between 10 and 12 dpi presented intense epileptiform cortical activity, beginning randomly after recording sessions started and sometimes followed a period of apparently normal recording (Fig. 2C). This epileptiform activity started with the presence of spiking activity that sometimes was more evident in the left or right hemispheres (Fig. 2D). Some minutes later, this activity became bilaterally synchronized and was characterized by the presence of polispike-waves or fast sharp waves that could last up to 7 minutes (Fig. 2E-G). The interspersed occurrence of depressed activity with the epileptic activity was progressively observed in the following minutes (Fig. 2H-J) and could culminate with the interruption of the epileptiform activity (Fig. 2K). During the 2 h recording session performed daily for every animal, they all (n=4) presented at least two of these epileptiform episodes. Behavioral changes observed during the occurrence of these epileptiform activities included freezing, tremor and tail erection. During the recording depression that followed each of the epileptic episodes, animals were agitated with increased exploration behavior for 1-2 min. No sign of epileptiform activity were observed in daily 2h-long recording sessions performed in mock- or iZIKV-injected mice, which showed mainly alternating fast and slow waves of low amplitude (Fig. 2B and Suppl. Fig. 1). Altogether, these results establish that our experimental model recapitulates spontaneous self-resolving seizures during childhood that are a frequent consequence of ZIKV infection. As neonatal ZIKV-infected mice were followed out of the acute phase of viral replication, seizure episodes decreased until they became undetectable in daily recording sessions performed with adult mice (∼100 dpi, see Fig. 2A) and no sign of electrographic seizures was detected in one ZIKV-infected ∼90 dpi mouse.

Then, we further examined the susceptibility of ZIKV-infected mice to develop drug-induced seizures after administration of pentylenetetrazol (PTZ), a GABA receptor antagonist widely used as a proconvulsant agent in animal models *(39)*. Interestingly, we found that both male and female adult (∼110 dpi) mice that had been infected with ZIKV at postnatal day 3 showed decreased latencies to the first seizure (Fig. 2L, N) and presented more seizure episodes (Fig. 2M, O) after PTZ treatment than mock-injected mice. The increased sensitivity to pharmacologically-induced seizures in adult mice indicates that early-life ZIKV infection leads to long-term neurochemical imbalances beyond the acute phase of viral replication. The results further suggest that the increased susceptibility to seizures may be yet another outcome in human adults that were exposed to ZIKV during the perinatal period.

### Neonatal ZIKV infection induces persistent motor and cognitive dysfunction in mice

Hypertonia, dystonic movement, arthrogryposis, difficulty of voluntary movement and grasp reflex are among the persistent motor dysfunctions seen in normocephalic babies congenitally exposed to ZIKV *(9)*. Hence, we next investigated the impact of early-life ZIKV exposure on the reflexes and behavior of young and adult mice. Since several early-life stressors may have differential impacts on subjects of both sexes *(40, 41)*, we evaluated separately the impact of ZIKV infection on male and female mice. To initially determine whether ZIKV infection was associated with a delay in development of neonatal reflexes, mice were evaluated daily in different reflex tests early after infection. No differences were found between ZIKV- and mock-injected pups in time to turn on the negative geotaxis (Suppl. Fig. 3A, B) and righting reflex (Suppl. Fig. 3C, D) tests, as well as in the grasping reflex score (Suppl. Fig. 3E, F). However, daily visual assessments suggested that mice showed a progressive increase in motor dysfunction, particularly involving the hind limbs, as ZIKV viral load increased in the brains. Therefore, ZIKV-infected male and female mice were tested in the Hind-Limb Suspension (HLS) test at 9 dpi, and in the Pole Test between 15 and 21 dpi. Both male (Fig. 3A) and female (Fig. 3B) ZIKV-infected mice showed decreased muscle strength compared to mock-injected animals in the HLS test. Male and female mice also showed motor dysfunction in the Pole Test, which is frequently used to evaluate fine motor function in young to adult rodents *(42)*: ZIKV-infected mice showed increased latencies to orient downwards (Fig. 3C,E) and to descend the extent of the pole (Fig. 3D,F).

**Figure 3.**
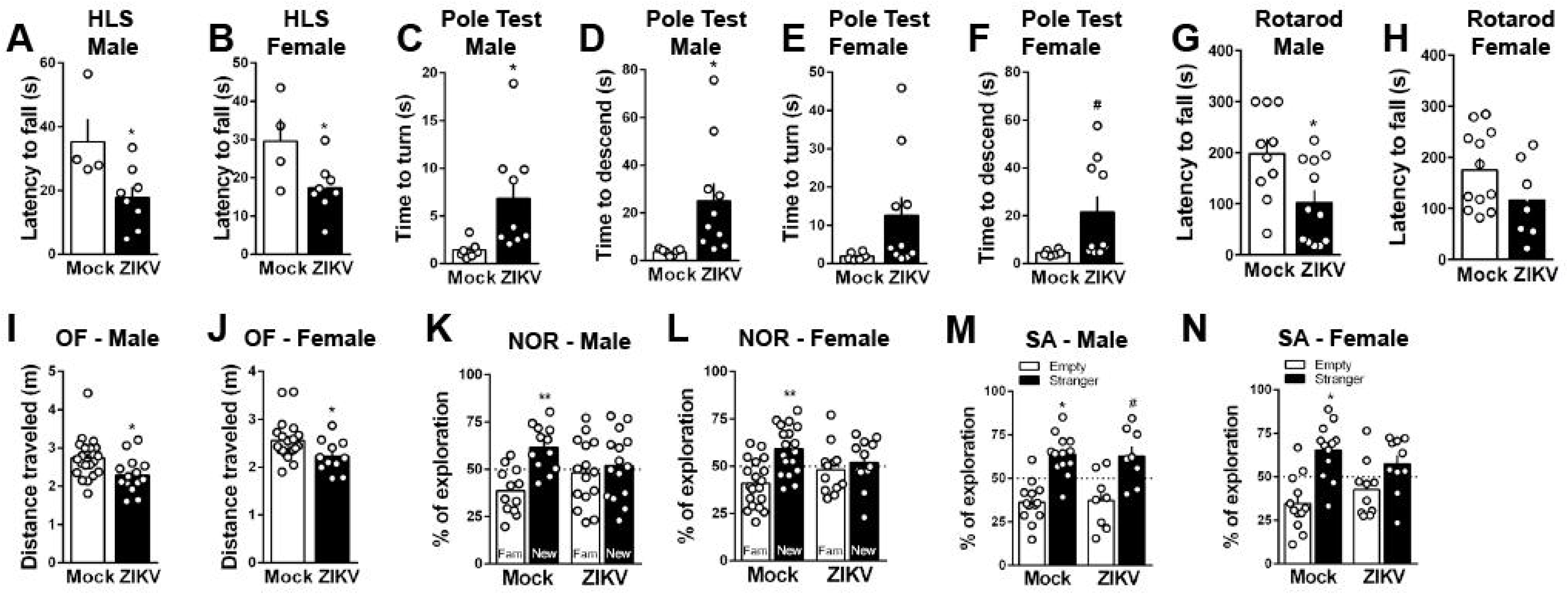
Neonatal ZIKV infection induces persistent motor and cognitive dysfunction in mice. Mice were infected by subcutaneous injection of 10^6^ PFU ZIKV or mock medium at post-natal day 3. (A, B) Latency to fall in the hind limb suspension test performed in male (A) and female (B) mice at 9 dpi. (C-F) Time to turn and time to descend in the pole test performed in male (C, D) and female (E, F) mice at 15-21 dpi. (G, H) Latency to fall in the rotarod for male (G) and female (H) mice at ∼90 dpi. (I, J) Distance traveled in the open field test by male (I) and female (J) mice at ∼90 dpi. (K, L) Percentage of exploration towards familiar (Fam.) and novel (New) objects in the novel object recognition test performed in male (K) and female (L) mice at ∼90 dpi. (M, N) Percentage of exploration of empty cage or cage containing a stranger mouse in the social approach task for male (M) and female (N) mice at ∼100 dpi. Data are expressed as mean ± S.E.M. HLS = hindlimb suspension test; OF = open field test; NOR = novel object recognition test; SA = social approach test. In **A**: *p = 0.029, in **B**: *p = 0.0421, in **C**: *p = 0.0243, in **D**: *p = 0.0287, in **F**: #p = 0.0662, in **G**: *p = 0.0149, in **I**: *p = 0.0164, in **J**: *p = 0.0322, Student’s *t* test. In **K**: **p = 0.0068, in **L**: **p = 0.0049, in **M**: *p = 0.001 and #p = 0.0569; in **N**: *p = 0.0064, one-sample Student’s *t* test compared to fixed value 50.

Infections of the central nervous system often lead to significant morbidity and persistent disability *(43)*, and the long-term effects of ZIKV infections have never been directly addressed. To determine whether this effect translated into motor and locomotor alterations in adulthood, animals were subjected to the rotarod and open field tasks (∼90 dpi). While only male ZIKV-infected mice showed a statistically significant smaller latency to fall in the rotarod paradigm compared to mock-injected groups (Fig. 3G, H), mice from both sexes showed impaired locomotor behavior, revealed by a decrease in total distance travelled in the open field task (Fig. 3I, J). We also examined whether early-life exposure to ZIKV, besides causing locomotor damage, also induced cognitive dysfunction in the adult life. To this end, adult mice infected with ZIKV at post-natal day 3 were trained and tested in the novel object recognition (NOR) paradigm (at ∼90 dpi). Both male (Fig. 3K) and female (Fig. 3L) infected with ZIKV failed the NOR task, indicating memory impairment in adult life.

Epidemiological and experimental evidence link perinatal infection to the development of neuropsychiatric diseases, including schizophrenia and autism *(44)*, disorders that have as a common feature impaired social interaction *(45, 46)*. This prompted us to evaluate whether early-life ZIKV infection affects social interaction in adult mice (∼100 dpi) employing the three-chamber social approach test (Fig. 3M, N). We found that, for male mice, ZIKV infection did not affect preference for exploring the stranger mouse over the empty chamber (Fig. 3M), while female ZIKV-infected mice showed impaired sociability, as demonstrated by similar exploration times of the empty and stranger mouse cages (Fig. 3N).

### Neonatal ZIKV exposure induces increased oxidative stress, causing brain inflammation and persistent neuropathological alterations in mice

In order to explore the pathological basis for the persistent consequences of early life ZIKV exposure, we performed neuropathological examination of the brains of mice shortly after seizures ceased (23 dpi). Severe brain atrophy with ventriculomegaly was found in some animals (Fig. 4A). We also observed that the brains of ZIKV-infected mice showed multiple necrotic areas, predominantly in the hippocampus, thalamus, striatum and cortex (Fig. 4B-D). Necrosis, dystrophic calcifications with mineral deposition, apoptotic bodies, infiltration of inflammatory cells, in addition to a complete disruption of the normal cytoarchitecture in all hippocampal subregions (Fig. 4F-H) were also seen in brains of infected animals. Along with the intense neuroinflammatory profile, brains of ZIKV-infected mice frequently showed frequent perivascular lymphocytic cuffings (Fig. 4I-J). Interestingly, histological alterations seen in young ZIKV-exposed mice persisted in adulthood, as a widespread neuronal necrosis with more advanced mineral deposition, especially in the hippocampus, was observed in the brains of 100-days old ZIKV-infected mice (Fig. 4K-N). Additionally, inflammatory cells and shrunken basophilic neurons, also called ferruginated neurons, were frequently seen in brains of mice exposed to ZIKV in the neonatal period (Fig. 4N).

**Figure 4.**
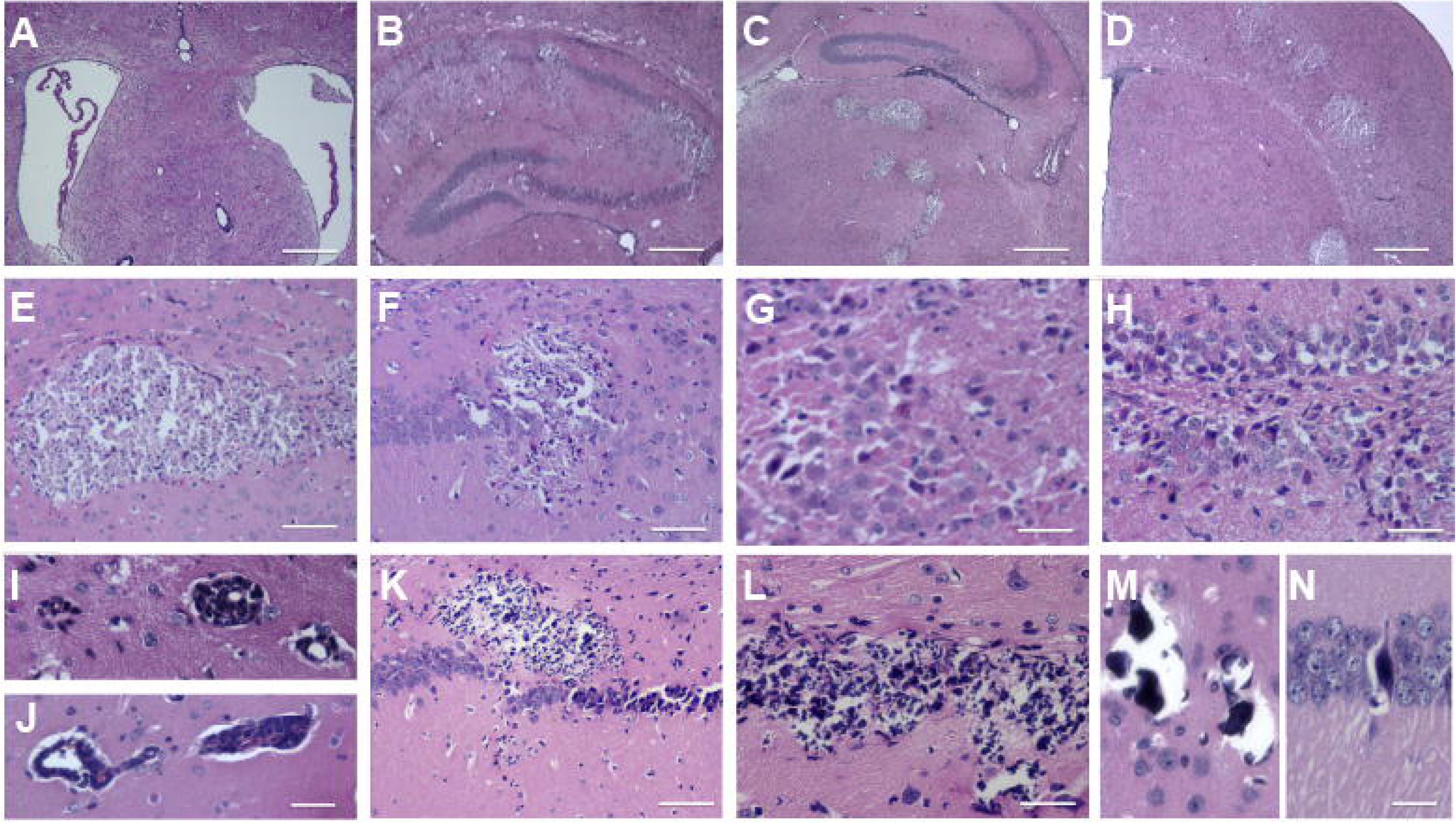
Neonatal ZIKV exposure causes severe and persistent neuropathological alterations in brains of mice. Mice were infected by subcutaneous injection of 10^6^ PFU ZIKV or mock medium at post-natal day 3. At 23 (A-J) or 100 dpi (K-N), mice were perfused, brains were dissected, processed and stained with hematoxylin and eosin. (A) Representative image of ventriculomegaly seen in brains of ZIKV-infected mice. (B-D) Representative images of multiple necrotic areas, predominantly in the hippocampus (B), thalamus (C), striatum and cortex (D) of ZIKV-infected mice. (E, F) Higher magnification representative images of dystrophic calcifications with mineral deposition, apoptotic bodies and inflammatory cells in brains of ZIKV-infected mice. (F-H) Representative images of complete disruption of normal cytoarchitecture in the CA1 (F), CA3 (G) and dentate gyrus (H) hippocampal subregions of ZIKV-infected mice. (I, J) Representative images of perivascular cuffings, seen in brains of ZIKV- infected mice. (K-M) Representative images of widespread neuronal necrosis with more advanced mineral deposition seen in brains of ZIKV-infected mice. (N) Representative image of a shrunken basophilic neurons (ferruginated neuron) seen in brains of ZIKV- infected mice. Scale bars = 450 μm (A, B), 300 μm (C, D), 50 μm (E, F, K, L), 35 μm (G- J), and 25 μm (M, N).

Neuroinflammation with microglial activation and production of proinflammatory cytokines in the brain plays an active role in epileptic disorders *(47)*. Thus, we performed immunohistochemistry in 23 dpi brain slices from mock and ZIKV-infected mice to evaluate astrogliosis. Compared to brains of mock-infused mice, we found that brains of infected mice showed markedly increased immunoreactivity for GFAP (Fig. 5A-D) and Iba-1 (Fig. 5E-H), notably in the proximity of lesions and in the perivascular areas, and especially in the CA1 and dentate gyrus (DG) hippocampal regions. In addition, the peak of viral replication in the brain was accompanied by increased brain mRNA levels of several pro-inflammatory cytokines, including TNF-α, IL-6, IL-1β, and KC, in ZIKV-infected animals when compared to mock-injected mice (Fig. 5I-L). Brain cytokine expression did not increase in brains of mice injected with UV-inactivated ZIKV (iZIKV) at 12 dpi (Fig. 5I-L), indicating that increased expression of these pro-inflammatory mediators requires active viral replication and not simply the presence of viral particles.

**Figure 5.**
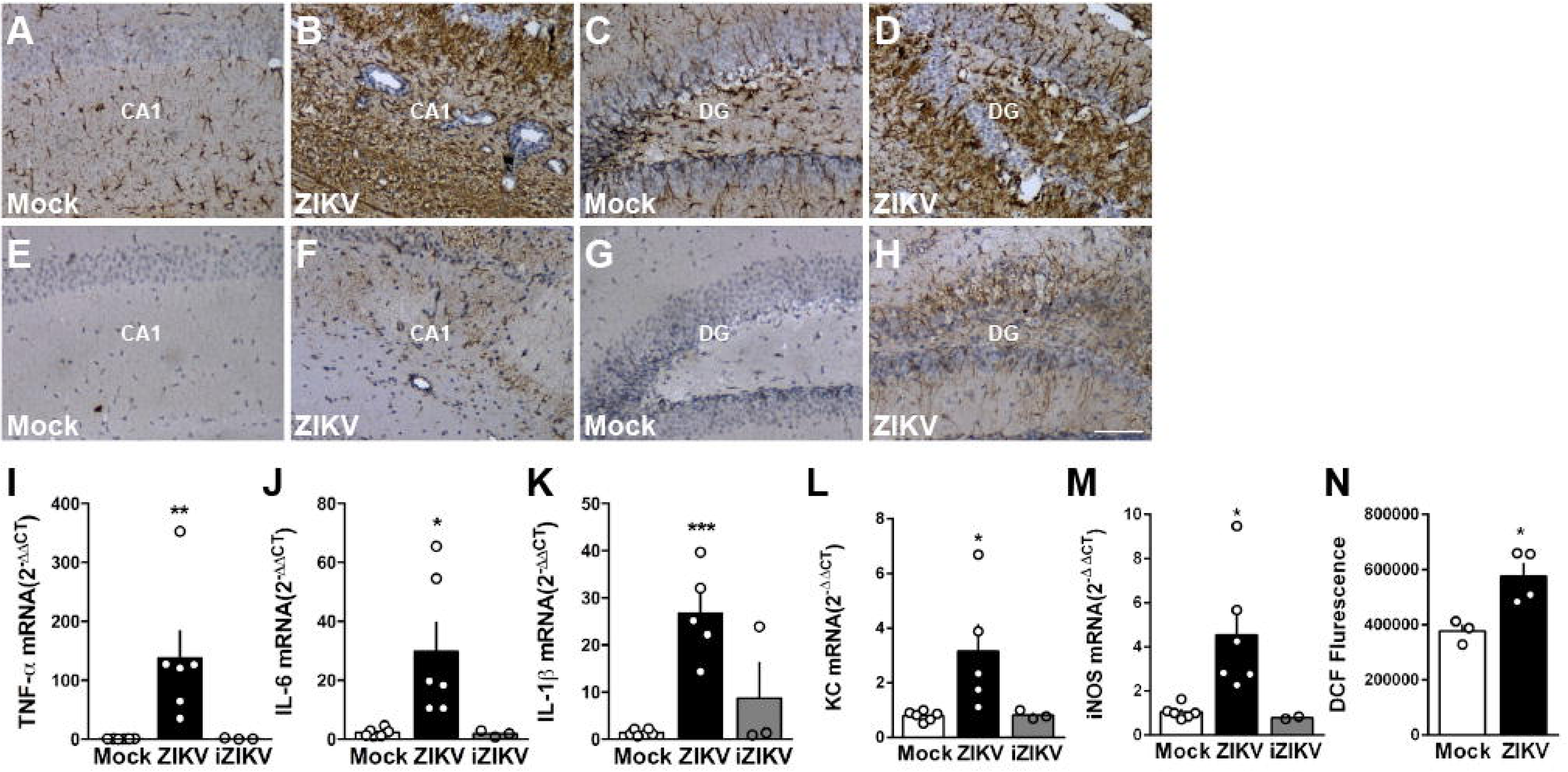
Neonatal ZIKV exposure induces brain inflammation and increased oxidative stress in mice. Mice were infected by subcutaneous injection of 10^6^ PFU ZIKV or mock medium at post-natal day 3. (A-H) At 23 dpi, mice were euthanized and brains were processed for immunohistochemistry. Representative images of brain sections immunolabeled for GFAP (A-D) and Iba-1 (E-H) show labeling on the CA1 and dentate gyrus areas of the hippocampus. Scale bar = 50 μm. (I-N) At 12 dpi, brains were dissected and processed for determination of mRNA levels of the following pro-inflammatory mediators by qPCR: (I) TNF-α, (J) IL-6, (K) IL-1β, (L) KC and (M) iNOS, or for determination of levels of reactive oxygen species (ROS). Data are expressed as mean ± S.E.M. In **I:** **p = 0.052, in **J**: *p = 0.0236, in **K**: ***p =0.0008, in **L**: *p = 0.0361, in **M**: *p = 0.0187, one-way ANOVA followed by Tukey. In **N**: *p = 0.0195 in Student’s *t* test.

**Figure 6.**
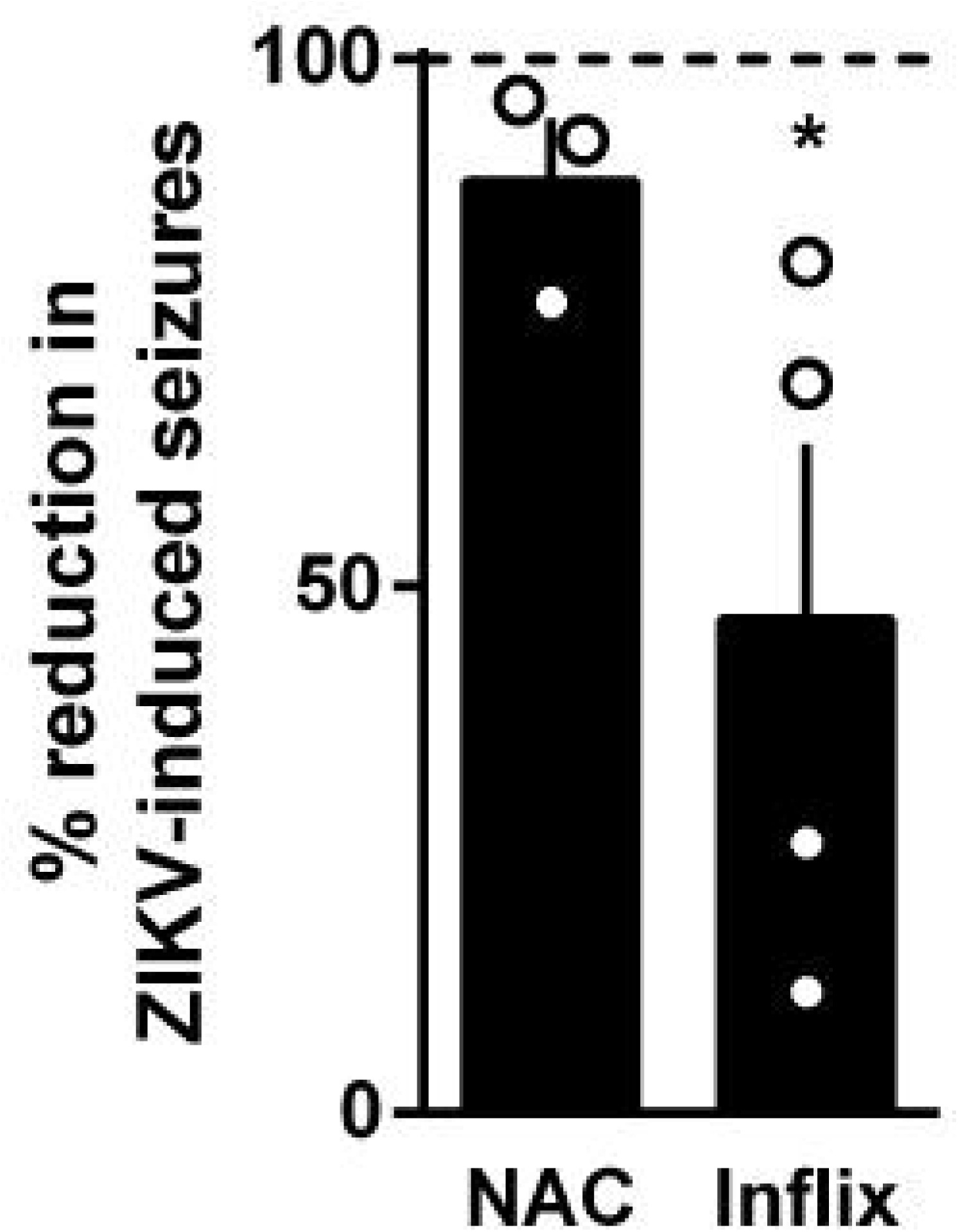
Infliximab, a TNF-α neutralizing antibody, prevents ZIKV-induced seizures in young mice. Mice were infected by subcutaneous injection of 10^6^ PFU ZIKV or mock medium at post-natal day 3 and were treated with daily ip injections of PBS, N-acetylcysteine (50 mg/kg) or Infliximab (20 μg/day) until 12 dpi. Percentage of animals with seizures in each litter was obtained from one hour-long recording session performed at 12 dpi in ZIKV-infected mice. Data are expressed as mean ± S.E.M. and normalized to percentage of animals showing seizures observed in ZIKV-infected mice. #p = 0.0925 in Student’s *t* test compared to fixed value 100.

Excessive generation of reactive oxygen species (ROS) is an important mechanism underlying brain dysfunction due to the infection by other viruses *(48)* and oxidative stress has been implicated in the pathology of acute seizures and epilepsy *(49)*. Accordingly, we found that the expression of the inducible form of nitric oxide synthase (iNOS) was increased in the brains of ZIKV-infected mice compared to mock animals at 12 dpi (Fig. 5M). Since nitric oxide (NO) can account for an increased generation of ROS *(50)*, we then hypothesized whether ZIKV replication in the brain promoted an increase in ROS generation. In agreement with our hypothesis, we found an increased fluorescence of the DCF dye in brain extracts from ZIKV-infected mice compared to mock mice (Fig. 5N), indicating a higher rate of cerebral ROS production associated with ZIKV infection.

### Infliximab, a TNF-α neutralizing antibody, prevents ZIKV-induced seizures in young mice

Both oxidative stress and inflammation have been shown to be sufficient to induce seizures in different animal models *(51, 52)*, and pharmacological approaches that combat both of them were shown to reduce epileptogenesis and lower seizure frequency. In order to investigate whether decreasing brain levels of ROS or blocking brain inflammation in ZIKV-infected mice prevented seizures, pups were treated with the antioxidant N-acetylcysteine (NAC) or the TNF-α neutralizing monoclonal antibody infliximab from the day of infection (P3) until seizure evaluation at 12 dpi. While NAC-treated litters showed only a mild reduction in the percentage of mice showing seizures, infliximab-treated litters showed a significant decreased in the percentage of mice showing seizures. These findings suggest that targeting inflammation during the acute phase of ZIKV infection is a useful approach to prevent development of seizures and possibly also the long-term behavioral consequences of this early life infection.

## Discussion

ZIKV infection during pregnancy has been associated to serious neurological conditions in fetuses, including microcephaly and brain malformations *(17)*. However, this severe clinical observation appears to be the most extreme consequence of congenital ZIKV exposure, and a spectrum of abnormalities and dysfunctions are expected to emerge. Since the long-term consequences of ZIKV infection are still unknown and have never been directly addressed, it is possible that the true magnitude of ZIKV epidemics is currently largely underestimated.

The negative impact of ZIKV to the developing central nervous system have been evaluated in immunodeficient mice or human brain organoids *in vitro (18–21, 53)*. These models show limited application for evaluation of the host immune response and of virus-induced behavioral alterations. Here, we described a model of neonatal ZIKV infection in immunocompetent mice to assess the acute and late behavioral and neuropathological consequences of early life virus exposure. It is important to notice that our model has a strong translational potential with fetal exposure, since this period of rodent brain development resembles the last trimester of gestation in humans *(33)*. Using this mouse model, we here show that neonatal ZIKV peripheral inoculation in mice leads to a ∼40% death rate and a persistent reduction in body weight gain until adulthood (∼69 dpi mice), suggesting that there are persistent effects in the general development of infected individuals associated to viral exposure. Studies have reported that normocephalic babies congenitally exposed to ZIKV show head growth deceleration beginning several months after birth, which results in postnatal-onset microcephaly *(11)*. In agreement, we here found that mice submitted to early-life ZIKV s.c. inoculation show macroscopic brain atrophy late (23 dpi) after infection.

Previous studies have detected ZIKV in the brains of neonatal mice shortly (5-10 dpi) after systemic inoculation *(22, 54)*. In agreement, we found that ZIKV subcutaneously inoculated successfully reaches and replicates in the mouse brain, with a replication peak between 6 and 12 dpi. Surprisingly, in our model ZIKV RNA was still detectable in adult mice (∼100 dpi) submitted to early-life viral infection. Persistent replication of the virus was confirmed by the presence of negative RNA strand in the brains of mice at this late time point, which represents an important long-term observation considering the lifespan of rodents. These findings bring clinically relevant evidence that brain replication of ZIKV might persist longer after the acute phase of disease.

Seizures are commonly reported consequences of infection by several viruses *(55)*. In particular, seizures have been reported following ZIKV infection in adult patients *(56)* and studies have shown that 50-62% of microcephalic babies due to ZIKV congenital infection develop this clinical manifestation *(10, 16)*. A variety of changes in sleep EEG patterns were found in these babies, with a predominance of interictal epileptogenic activity and hypsarrhythmia, and in other cases electrographic seizures were identified even in the absence of clinical manifestation *(10, 16)*. In addition, follow-up studies performed with normocephalic babies exposed *in uterus* to ZIKV suggest that seizures and epilepsy are clinical manifestations seen in ∼50% of babies *(11)*. In agreement with these clinical evidences, we observed a high incidence of seizures in neonatal mice exposed to ZIKV. Thus, we perform temporal analysis of seizure development and the electrographic alterations associated to brain viral replication. Seizures were detected from 9 dpi, and reached higher incidence at 12 dpi, which coincided with the peak of viral RNA detected in the brain. This finding is in agreement with a preliminary report, in which authors performed a primary visual assessment in mice following ZIKV infection and reported self-resolving seizures *(24)*. Here, we further showed that cortical electroencephalographic recordings from ZIKV-infected mice showed severe and recurrent epileptiform activity between 10-12 dpi, with the presence of spiking activity, polispike waves and fast sharp waves. To the best of our knowledge this comprises the first characterization of ZIKV-induced seizures in mice. Our data also show that this behavioral outcome in the acute phase of infection is dependent on high levels of viral replication in the brain, since seizures were almost entirely absent in mice as soon as ZIKV or UV-inactivated ZIKV was infused. We followed mice post the acute phase of infection and found that the number of seizure episodes were much smaller at 18 dpi and were not observed in adult (∼100 dpi) mice. These findings show that our model is useful in reproducing the epileptic effect of ZIKV in infants and that spontaneous seizures associated to viral exposure might be reversible as patients grow older. However, our results also establish that adult mice exposed to ZIKV during development show increased susceptibility to chemically-induced seizures and a higher number of seizure episodes following chemical stimulation, showing that the neurochemical imbalances induced by viral infection are long lasting. Studies have demonstrated that several viral infections that affect the central nervous system may induce seizures both in experimental models and in humans *(57–59)* during the acute phase of infections *(60)*. Epidemiological analyses have also demonstrated that these viral infections often lead to development of persistent epilepsy after infection is resolved *(61–64)*. Galic and colleagues (2009) showed that neonatal exposure to poli(I:C) increased susceptibility to seizures and memory deficits in adult rats, and that these behavioral changes were associated to neuroinflammation and an increased hippocampal expression of NMDA receptors *(65)*. The results reported in our study indicate that even normocephalic ZIKV-exposed infants might be at increased risk of being diagnosed with epilepsy, and further suggest that a long-term follow-up of exposed patients should be pursued by public health services, especially in countries affected during the 2015 epidemics.

A number of motor alterations have been reported in babies exposed to ZIKV *in uterus*, including hypertonia, hemiparesis, dyskinesia/dystonia and arthrogryposis, among several others *(9)*. Moreover, ZIKV has been shown to infect peripheral neurons, causing cell death *(66)*. Here, neonatal mice infected with ZIKV were submitted to different tasks to evaluate neonatal reflexes, neuromuscular function, muscle strength and locomotor behavior throughout development. ZIKV-injected pups showed normal performance in reflex evaluation performed early after infection. These results are in contrast with a previous report of impaired performance in the righting reflex induced by an African ZIKV strain *(54)*, which might be explained by the fact that this strain is much more aggressive to the murine nervous system compared to the strain used in our study *(67)*. As we continued to follow the behavior of ZIKV-infected mice, we found that they showed poor performance in the hindlimb suspension test (9 dpi), pole test (15-21 dpi), rotarod and open field test (∼90 dpi), suggesting that impaired motor function is a consistent and long-lasting effect of infection. Early life exposure to several viruses has been associated with development of learning problems *(68, 69)* and neuropsychiatric disorders *(70)* later in life. Here, we report that male and female adult mice infected with ZIKV at post-natal day 3 show impaired performance in the novel object recognition (NOR) task (∼90 dpi), a non-rewarded, ethologically relevant paradigm widely used to assess declarative memory in rodents *(71)*. Similar results have recently been reported using two different tasks to evaluate spatial and aversive memories *(54)*. These results strongly suggest that ZIKV-exposed infants might develop learning problems, which can seriously compromise patient’s life quality. Impaired sociability is a hallmark feature of several neuropsychiatric disorders including autism. Here we report that ZIKV-infected female, but not male, mice showed poor performance in the social approach (SA) task (∼100 dpi). These results suggest neuropsychiatric disorders might be a long-term consequence of perinatal ZIKV exposure and that additional epidemiological studies should address this subject in more detail.

To date, neuropathological studies described a spectrum of changes associated with congenital ZIKV infection, including calcification, necrosis, nerve cell degeneration, microgliosis, hypoplasia, and ventriculomegaly in fetal and newborn brain tissue *(72–75)*. Here we found that brains of mice infected with ZIKV during neonatal period show macroscopic brain atrophy and several histologic cerebral lesions, many of which persistent until animals reach adulthood. We report multifocal areas of necrosis in several brain regions, especially in the hippocampus, and show that ZIKV-induced necrosis is often composed by calcifications and apoptotic cells. Intense inflammatory infiltrations are widespread throughout the brains, with large presence of perivascular cuffings and microglial nodes, which are hallmarks of viral encephalitis *(76, 77)*. Brains of patients or animals that exhibit epilepsy (eg. adult pilocarpine-induced seizure) show intense microgliosis and massive neuronal loss in CA1, CA3, and DG hippocampal subregions *(78)*. Despite this similar vulnerability of hippocampus to damage, the histopathological pattern of brain damage in ZIKV-infected mice found in our study is distinct from pilocarpine-induced lesions. This histological pattern suggests that hippocampal lesions induced by ZIKV infection results in seizure, opposing to the pilocarpine model in which epileptic crises result in hippocampal lesions. The spectrum of neuropathological findings observed in our model is, in several aspects, analogous to those seen in fetuses and newborns with ZIKV-congenital syndrome *(75)*.

Brain inflammation has been increasingly implicated in the pathogenesis of neurodevelopmental diseases and in the long-term consequences of aversive early life experiences *(79)*. Increased levels of pro-inflammatory cytokines have been reported in placentas and in brains of newborns exposed to ZIKV *(13)*. Here, we showed that viral replication was accompanied by increased brain expression of several proinflammatory immune mediators. Immunohistochemical analyses confirmed the presence of an exacerbated and aberrant microglial activity in the brains of neonatal ZIKV-infected mice. We also showed that blocking the effects of TNF-α starting shortly and persistently after ZIKV-infection, we were able to significantly reduce the occurence of seizure in young mice. On the other hand, although we also found ZIKV-infection induced increased ROS generation in brains of mice, treatment with the antioxidant NAC was not efficient in preventing seizure episodes. These results suggest that inflammation is a key mechanism underlying ZIKV-induced seizures and that patients exposed to the virus during the perinatal period could benefit from the use of anti-inflammatory strategies.

In conclusion, our findings comprise the first experimental evidence that ZIKV perinatal exposure is associated to long-term deleterious neuropathological and behavioral consequences. A wide spectrum of neurological and neuropsychiatric manifestations might be expected as a consequence of perinatal ZIKV exposure, especially as an outcome of the 2015 epidemics, which affected a large number of individuals across the Americas. Moreover, our data suggest that targeting inflammation during the acute phase of infection might be a useful approach to prevent ZIKV-induced seizures. It is now clear that simply monitoring the birth prevalence of congenital microcephaly is an insufficient device to measure the burden of ZIKV neuropathology in exposed children and adolescents.

## Acknowledgments

This work was supported by grants from Brazilian funding agencies: Fundação de Amparo à Pesquisa do Estado do Rio de Janeiro (FGDF, STF, ATP, IAM., CPF and JRC), Fundação de Amparo à Pesquisa do Estado de São Paulo (EAC), Institutos Nacionais de Pesquisa - Instituto Nacional de Neurociência Translacional (FGDF, EAC, STF), Conselho Nacional de Desenvolvimento Científico e Tecnológico (PSF, GN, FGDF, EAC, STF, ATP, CPF, IAM and JRC), Institutos Nacionais de Pesquisa - Inovação em Medicamentos e Identificação de Novos Alvos Terapêuticos (CPF), Coordenação de Aperfeiçoamento de Pessoal de Nível Superior (INOS, JVF, LF, DJLLP), and Financiadora de Estudos e Projetos (ATP). We thank Melissa Florence Marques Miécimo da Silva, Jadilma Araujo Ferreira for technical support, and Ana Claudia Rangel for competent lab and project management.

## Figure Legends

**Supplemental Figure 1. Schematic drawing of electrodes implanted in the right (R1-R2) and left (L1-L2) cortex of mice.** Mice were infected by subcutaneous injection of 10^6^ PFU ZIKV or mock medium at post-natal day 3. Electrographic recordings obtained in the right and left cortex from mock- (B-E) and UV-inactivated ZIKV-injected (F-I) mice (10 to 12 dpi). No sign of epileptiform activity is seen in neither groups. N = 2 mice/group.

**Supplemental Figure 2. Late viral brain replication and seizure evaluation in UV-inactivated ZIKV-injected mice.** Mice were infected by subcutaneous injection of 10^6^ PFU ZIKV or mock medium at post-natal day 3. (A) ZIKV-infected mice brains were dissected and processed for negative strand qPCR analysis at 12 or 100 dpi. (B) Percentage of mice with seizures in ZIKV and UV-inactivated ZIKV (iZIKV)-injected groups were evaluated in one hour-long recording sessions performed at 9, 12 and 18 dpi (n = 4 litter ZIKV and 3 litters iZIKV). Data are expressed as mean ± S.E.M.

**Supplemental Figure 3. Neonatal ZIKV infection is not associated to altered development of reflexes in newborn mice.** Mice were infected by subcutaneous injection of 10^6^ PFU ZIKV or mock medium at post-natal day 3, and negative geotaxis (A-B), righting reflex (C-D) and grasping reflex (E-F) were measured in male and female mice, respectively, until 7 dpi.

